# Physiologic Regulation of Lipid Oxidation and Ferroptosis By Vitamin E, High-Density Lipoprotein, and Scavenger Receptor Class B Type 1

**DOI:** 10.1101/2024.11.13.623420

**Authors:** Jonathan S. Rink, Andrea E. Calvert, Alexandra Moxley, Adam Yuh Lin, SonBinh T. Nguyen, Leo I Gordon, C. Shad Thaxton

**Affiliations:** Northwestern University, Feinberg School of Medicine, Department of Urology, Chicago, IL 60611; Northwestern University, Feinberg School of Medicine, Department of Medicine, Division of Hematology/ Oncology, Chicago, IL 60611; Northwestern University, Feinberg School of Medicine, Simpson Querrey Institute for BioNanotechnology, Chicago, IL 60611; Northwestern University, Feinberg School of Medicine, Robert H. Lurie Comprehensive Cancer Center, Chicago, IL 60611; Northwestern University, Weinberg College of Arts and Sciences, Department of Chemistry, Evanston, IL 60208

**Keywords:** Ferroptosis, α-tocopherol, high-density lipoproteins, scavenger receptor class B type 1

## Abstract

Ferroptosis is a form of cell death caused by oxidative (-OOH) damage to phospholipids (PL) that comprise the cell membrane. Understanding mechanisms of ferroptosis is important because of the role it plays in human disease, including cancer and neurodegeneration. Previous work has focused on the intracellular antioxidant enzyme glutathione peroxidase 4 (GPx4), which detoxifies PL-peroxides (PL-OOH) and prevents ferroptosis. Studies also show that alpha-tocopherol (α-toc), the active form of Vitamin E, is an exogenous lipophilic antioxidant that inhibits ferroptosis in cancer cells in vitro and in neurons and hematopoietic stem cells in vivo. The mechanisms by which α-toc engages cells to manipulate PL redox balance and ferroptosis are unknown. Lipoproteins, like high- and low-density lipoproteins (HDL and LDL), carry α-toc in the circulation, which compelled our investigation of their role in α-toc delivery and ferroptosis. Using cancer and neuronal cell models, here we show that lipoproteins, particularly HDL, deliver α-toc by binding the cell membrane receptor scavenger receptor class B type 1 (SR-B1) to prevent ferroptosis. Additionally, we synthesized SR-B1 targeted HDL-like particles that contained α-toc to validate data obtained with native lipoproteins. We reveal here a tunable cellular redox axis whereby native and synthetic HDLs containing α-toc target SR-B1 to reduce PL-OOH and prevent ferroptosis.

## INTRODUCTION

Ferroptosis is an iron-dependent cell death mechanism that results from shifting cellular redox balance to favor increased lipid reactive oxygen species (L-ROS) and the propagation of lipid oxidative damage to polyunsaturated fatty acid (PUFA) tail groups of cell membrane phospholipids (PLs)^1–3^. There is considerable interest in manipulating cellular PL redox balance and ferroptosis for next generation cancer therapies and to prevent neurodegeneration, among other applications^4,5^.

Endogenous intracellular proteins and pathways that regulate cell death by ferroptosis have been identified^1,6^. Importantly, acyl-CoA synthase long chain family member 4 (ACSL4) is the enzyme responsible for generating cell membrane PLs containing PUFA alkyl tail groups (PUFA-PL), which are targets for oxidation to PUFA-PL peroxides (PUFA-PL-OOH)^7,8^. Protecting the cell, GPx4 is a selenoenzyme that utilizes glutathione (GSH) as a co-factor to reduce PUFA-PL-OOHs to their less toxic alcohols (PUFA-PL-OHs)^9,10^. Finally, an intracellular lipophilic antioxidant, coenzyme Q10 (CoQ_10_), and its enzymatic redox partner, ferroptosis suppressor protein 1 (FSP1), can protect cell membrane PLs from oxidation^11^.

The active form of Vitamin E, alpha-tocopherol (α-toc), is an exogenous lipophilic antioxidant shown to localize to the cell membrane and reduce lipid oxidative damage^10,12,13^. Early in vitro studies that focused on pathways to induce oxidative cancer cell death by targeting the GSH/GPx4 axis revealed that α-toc rescues cells from cell death even when these protective pathways have been inhibited^14–16^. In *in vivo* animal models, α-toc has been shown to prevent oxidative cell death in multiple cell types, including neurons^13,17–19^ and hematopoietic stem cells^20^, even when the lipid protective functions of GPx4 have been reduced or genetically knocked out^17–20^.

Following dietary absorption in the gut, α-toc requires solubilization and transport in the blood stream by lipoproteins^21,22^. Specifically, low-density lipoprotein (LDL)^21,23,24^ and high-density lipoprotein (HDL)^21,25,26^ carry α-toc in the circulation for delivery to target cells. The LDL particles target the LDL receptor (LDLR) whereupon the LDL/LDL-R complex is internalized in a clathrin-dependent manner^27,28^. The LDL particle is then destined for degradation in the endolysosome whereby the degradation products are then distributed to the cell^27^. Studies using murine knockout models for LDLR show little difference in tissue α-toc content, suggesting that the role for LDLs in delivering α-toc is not rate limiting and that there are alternative mechanisms^26^. The high-affinity receptor for HDLs, SR-B1, was initially discovered as a receptor for acetylated LDL; however, it was later defined as a preferential receptor for HDL^29^. Lipid-rich HDLs bind SR-B1 and oligomerize SR-B1 at the cell membrane^30^. Upon binding, HDLs deliver and exchange lipids while docking at the cell membrane^30,31^. Studies in SR-B1 knockout mice reveal that the serum level of α-toc increases and that tissue levels of α-toc decrease in the gonads, brain and kidney^22,26,29,32,33^. Accordingly, HDLs are known to have antioxidant functions^34^, and an α-toc/HDL/SR-B1 axis is clearly involved in maintaining lipid redox balance in specific cells and tissues^26,35^.

To investigate α-toc, lipoproteins, and cell surface receptors as mediators of ferroptosis, we employed native HDL and LDL, including ones we manipulated to have higher α-toc content. Also, we developed a synthetic, organic-core templated HDL NP (ocHDL NP) with similar size, shape (spherical), charge (anionic) and surface chemistry [e.g., apolipoprotein A-I (apoA-I) and phospholipids] as native HDLs that bind to SR-B1^36^. The ocHDL NPs enabled us to control particle composition including α-toc content. Using native HDL and LDL, native HDL and LDL with increased α-toc, and α-toc loaded ocHDL NPs (α-toc ocHDL NPs), we find that α-toc delivery through SR-B1, but not through the LDL receptor (LDLR), mediates cell membrane lipid oxidation and inhibits ferroptosis in cancer cells and in neuronal cells. These data delineate an α-toc/HDL/SR-B1 axis of cellular protection against cell membrane lipid peroxidation. Furthermore, and for the first time, we reveal that targeting SR-B1 using native or synthetic SR-B1 ligands is a plausible and novel strategy to control PL oxidation and cell death via ferroptosis.

## RESULTS

### Native SR-B1 Ligands Rich in α-toc Rescue Cancer Cells from Ferroptosis Induced by The Small Molecule Covalent Inhibitor of GPx4, ML210

Diffuse large B-cell lymphoma (DLBCL) and clear cell renal cell carcinoma (ccRCC) cell lines were among the first shown to be particularly sensitive to ferroptosis after treatment with small molecule covalent inhibitors of GPx4, and cell death was rescued by addition of Vitamin E^10^. Our group demonstrated DLBCL sensitivity to ferroptosis using two lymphoma cell lines, Ramos and SUDHL4^37^. Thus, we explored α-toc delivery by lipoproteins and ferroptosis by employing Ramos, SUDHL4, and a ccRCC cell line (786-O). All three cell lines express SR-B1 (Figure S1A) and the LDLR (Figure S1B). We investigated whether native HDL or LDL could rescue the cancer cell lines from cell death when treated with a covalent inhibitor of GPx4, ML210^9^. First, we measured the amount of α-toc in native HDL and LDL particles per apolipoprotein [apolipoprotein A-I (apoA-I) for HDL and apolipoprotein B-100 (apoB-100) for LDL]. Native HDL particles had ∼1.0 ± 0.1 α-toc molecules per apoA-I, and LDL particles had 6.0 ± 0.03 α-toc molecules per apoB-100 (Table S1), consistent with previous reports^38–40^. To increase the α-toc content of both lipoproteins beyond what is observed physiologically, we incubated HDL and LDL with free α-toc followed by particle purification. The resultant α-toc lipoproteins (α-toc HDL and α-toc LDL) showed a moderate increase in α-toc to apolipoprotein ratios (1.6 ± 0.3 per apoA-I in α-toc HDL and 10.1 ± 0.1 per apoB-100 in α-toc LDL; Table S1).

We treated the Ramos, SUDHL4, and 786-O cancer cells with a dose of ML210 that induced >95% cell death in each cell line (250 nM for Ramos; 100 nM for SUDHL4; 1 µM for 786-O; Figure S2A-C). Rescue experiments were conducted by adding native HDL and LDL and HDL and LDL with increased α-toc (α-toc HDL and α-toc LDL; **Figure 1A-C**). We found that HDL rescued the cancer cells from ML210-induced cell death at lower concentrations compared with LDL (**Figure 1A-C**). The α-toc HDL and α-toc LDL demonstrated an enhanced ability to rescue ML210-induced cell death (**Figure 1A-C**). Despite ML210 being established as an agent that inhibits GPx4 and induces ferroptosis by increasing cell membrane lipid peroxidation, we employed the SUDHL4 cells to confirm that rescue from cell death was accompanied by a decrease in cell membrane lipid peroxide accumulation after treatment with the various lipoproteins. As expected, increased cell viability was accompanied by reduced cell membrane lipid oxidation measured using C11-BODIPY, indicating that ferroptosis was the dominant mechanism of cell death (**Figure 1D**).

**Figure 1:**
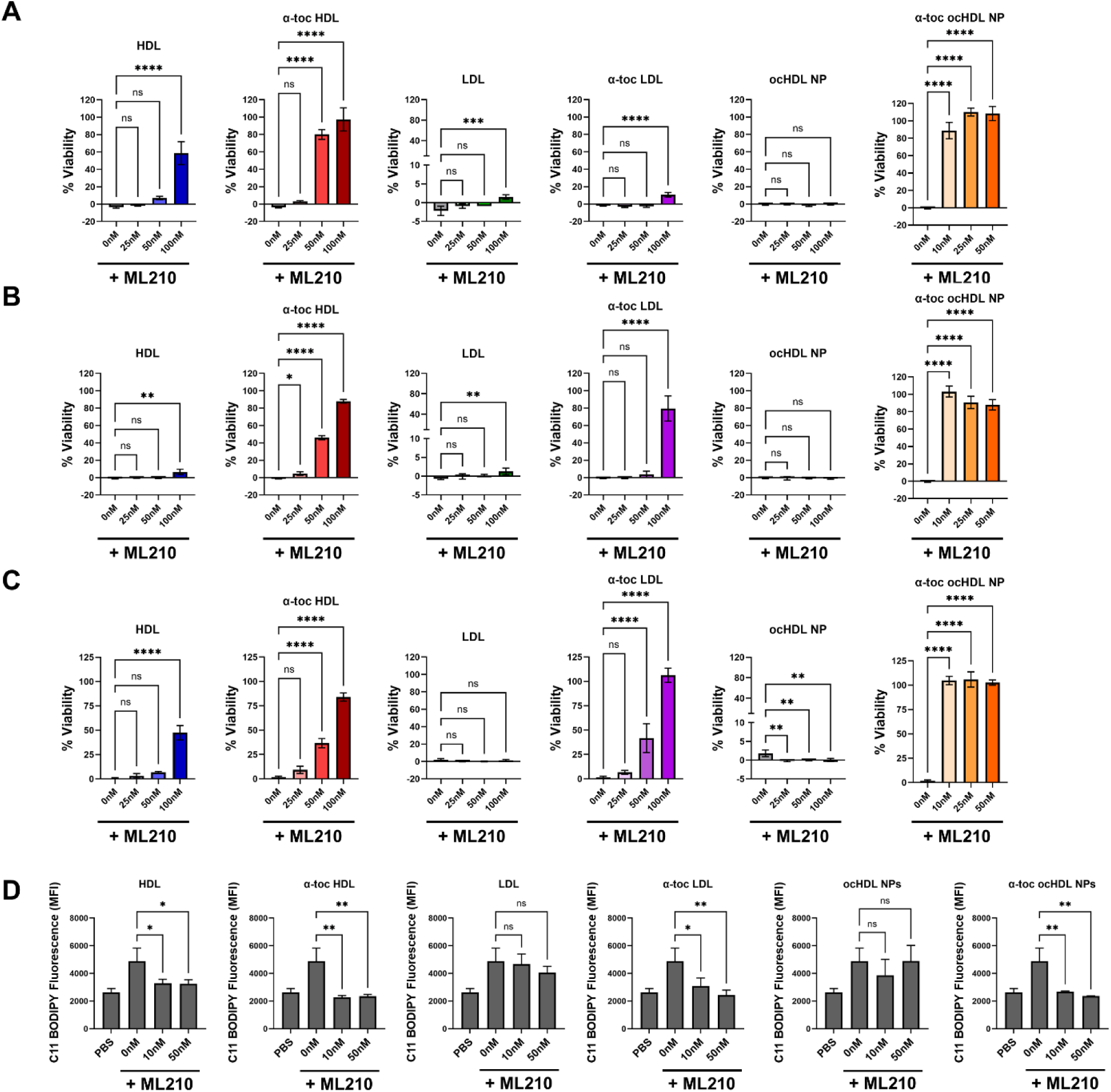
α-toc Delivery by Natural Lipoproteins Rescues ML210-Induced Cell Death in DLBCL and ccRCC Cells. A-C. Ramos (A), SUDHL4 (B) and 786-O (C) cells were treated with ML210 (Ramos – 250 nM, SUDHL4 – 100 nM; 786-O – 1 µM) in combination with increasing concentrations of HDL, α-toc HDL, LDL, α-toc LDL, ocHDL NP and α-toc ocHDL NP for 72 hrs. D. C11 BODIPY analysis of lipid peroxide accumulation in SUDHL4 cells treated with ML210 (100 nM) and HDL, α-toc HDL, LDL, α-toc LDL, ocHDL NP and α-toc ocHDL NP. Statistics: *p<0.05, **p<0.01, ***p<0.001 and ****p<0.0001 indicate statistical significance. No statistical difference (p>0.05) is noted by “ns.” See Supporting Information for more detailed statistical analyses.

### Native and α-toc HDLs effectively reduce cell membrane PL oxidation and rescue cancer cells from ferroptosis

Native HDLs are dynamic nanostructures and chemically heterogeneous^41–43^. Therefore, to isolate the effect of α-toc delivered by HDL, we utilized a synthetic HDL NP platform where particle composition could be controlled. In brief, this synthetic HDL NP platform utilizes an organic core (oc) molecule that serves as a core template to support the self-assembly of phospholipids and apoA-I into particles that we referred to as ocHDL NP^36^. The ocHDL NPs have similarities to native HDLs (e.g., apoA-I and phospholipids), including size, shape, negative surface charge, and ocHDL NPs target SR-B1^36^. We synthesized ocHDL NP with and without α-toc. The resultant α-toc ocHDL NPs had 3.5 ± 2.2 α-toc per apoA-I, which exceeds α-toc levels in natural HDLs or α-toc HDLs due to the increased core volume available for α-toc loading versus the natural cargo (e.g., cholesterol ester and triglycerides) known to occupy the core of native HDL (Table S1). The ocHDL NP and α-toc ocHDL NPs trended toward being larger than native HDL and α-toc HDL, which may have also contributed to higher α-toc loading, but the sizes were not statistically significantly different (Table S1). The α-toc ocHDL NPs did not differ in size compared with empty ocHDL NPs (p=0.9934; Table S1); however, the surface charge (zeta potential) of α-toc ocHDL NPs was significantly less negative than that of empty ocHDL NPs (Table S1). The less negative zeta potential in α-toc ocHDL NP may be due to packing of neutral α-toc in the surface and core lipids of the ocHDL NP or alterations in phospholipid packing and/ or apoA-I conformation upon α-toc loading^44,45^. We utilized the ocHDL NPs to investigate the delivery of α-toc and the effects of the materials on PL oxidation and ferroptosis.

Like in the previous experiments, we added increasing concentrations of ocHDL NPs or α-toc ocHDL NPs to the DLBCL and ccRCC cell lines treated with ML210. We found that ocHDL NPs without α-toc did not rescue the cell lines from ML210-induced cell death; however, addition of the α-toc ocHDL NPs led to a significant increase in cell survival despite ML210 treatment. (**Figure 1A-C**). Cell rescue was accompanied by a significant decrease in cell membrane lipid oxidation measured using C11-BODIPY (**Figure 1D**).

To confirm these findings using an alternative mechanism of inhibiting GSH/GPx4 we repeated these experiments using the ferroptosis inducer erastin instead of ML210. Erastin inhibits cellular uptake of the GSH precursor, cystine, through system x_c_ ^−^ to induce ferroptosis^46^. We found that both natural and synthetic lipoproteins loaded with α-toc were capable of rescuing erastin-induced ferroptosis (**Figure 2**).

**Figure 2:**
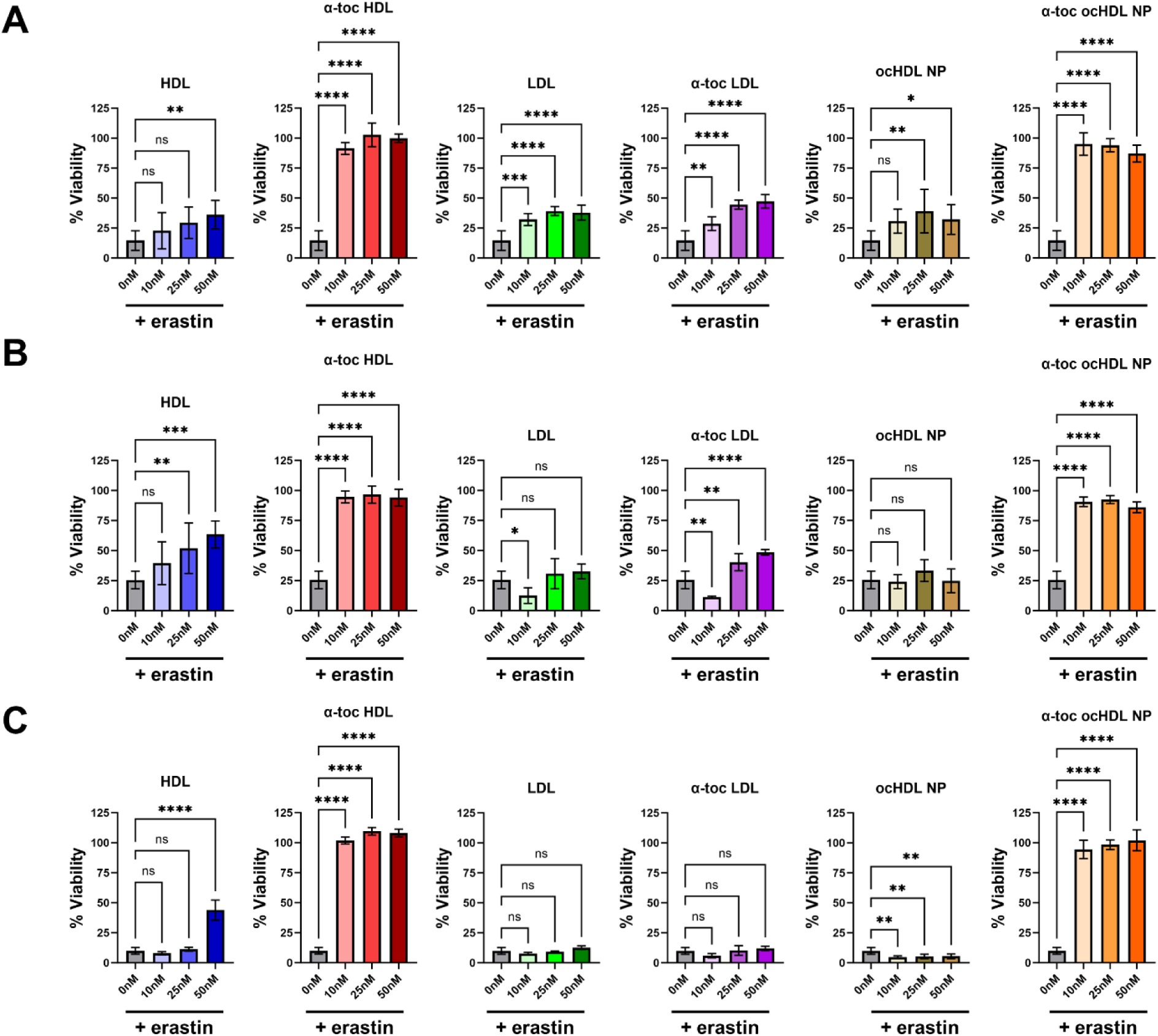
Lipoprotein Rescue of Erastin-Induced Cell Death. A-C. Rescue of Ramos (A), SUDHL4 (B) and 786-O (C) cells from erastin-induced cell death by HDL, α-toc HDL, LDL, α-toc LDL, ocHDL NPs and α-toc ocHDL NPs. Statistics: *p<0.05, **p<0.01, ***p<0.001 and ****p<0.0001 indicate statistical significance. No statistical difference (p>0.05) is noted by “ns.” See Supporting Information for more detailed statistical analyses.

### SR-B1, But Not LDLR, Mediates Uptake of α-toc From Lipoproteins

Both HDLs and LDLs are capable of binding SR-B1 and the LDLR^29,32^. Further, both receptors have been shown to be expressed in a number of different malignancies, including DLBCL and ccRCC^47,48^, and they have been implicated in central nervous system physiology and pathology, including neurodegeneration^49,50^. To assess the role of the individual receptors in α-toc delivery and ferroptosis, we used siRNA to reduce expression of SR-B1 and LDLR. We conducted these experiments in the adherent 786-O cells as they are more conducive to siRNA transfection than the DLBCL cell lines that both grow in suspension. Consistent with our previous data collected in ovarian cancer^51^, knockdown of SR-B1 led to decreased GPx4 expression (Figure S3). Decreased GPx4 expression correlated with increased PUFA-PL-OOH accumulation and increased cell death (Figure S3). Knockdown of the LDLR was confirmed by flow cytometry (**Figure 3A**). Reduction of the LDLR did not significantly change SR-B1 (**Figure 3A**) or GPx4 expression (Figure S4). Using the LDLR knockdown cells, we investigated whether α-toc containing HDL and LDLs were capable of rescuing ML210-induced ferroptosis. We demonstrated enhanced rescue in the siLDLR cells compared to wild type (Neg Control) cells and those treated with scrambled (SCR) siRNA (siSCR; **Figure 3B-D**). This effect could be due to increased binding of the LDLs to SR-B1, as previous studies have shown that LDLs bind to SR-B1^29,52^, or to increased transfer of α-toc from LDLs to HDLs that then deliver α-toc via binding SR-B1^22^. Regardless, these data indicate that SR-B1, rather than LDLR, is the critical receptor for α-toc delivery by lipoproteins.

**Figure 3:**
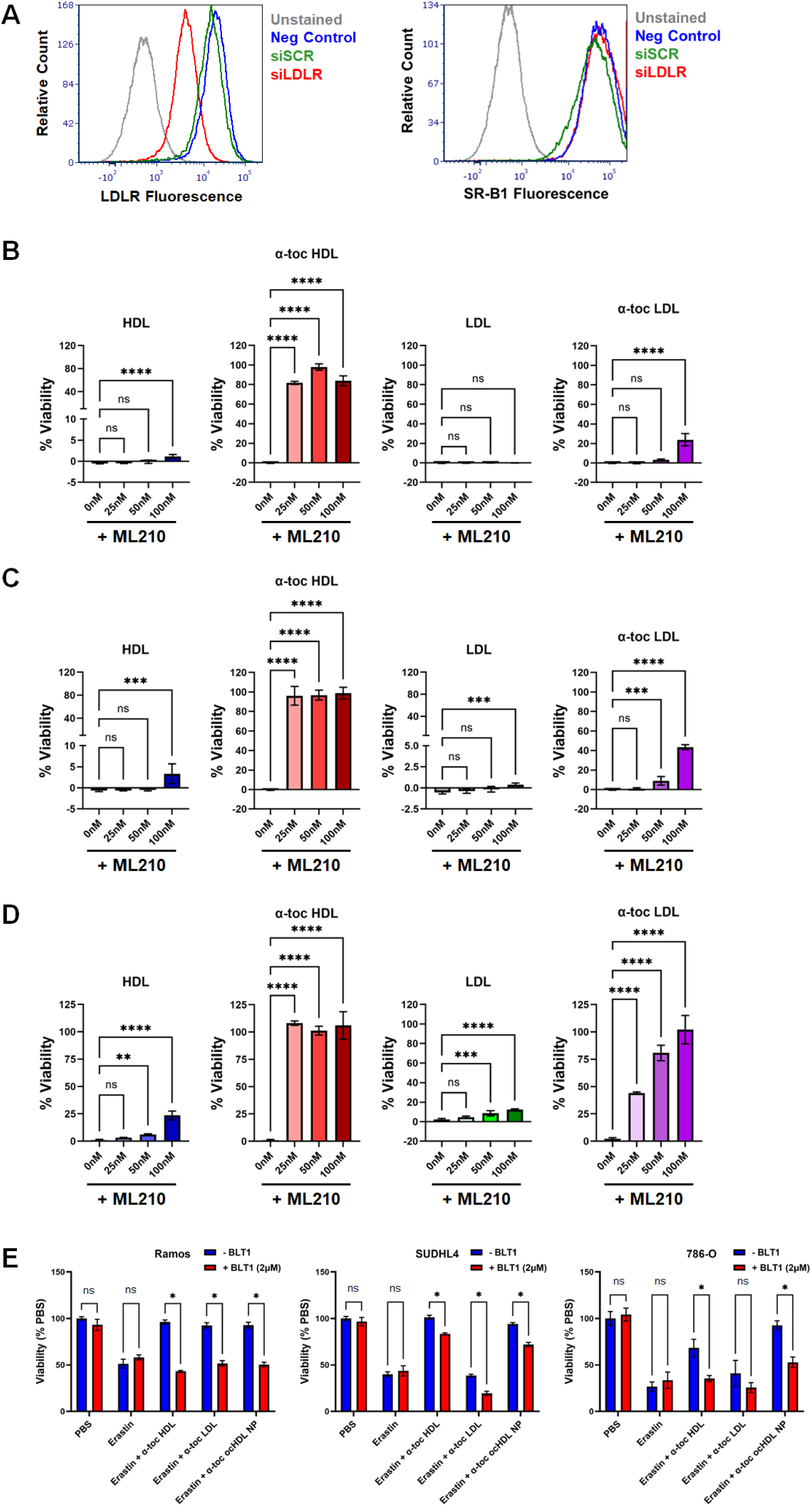
SR-B1, But Not LDLR, Mediates α-Toc Rescue in Cancer Cells. A. Flow cytometry analysis of LDLR (left) and SR-B1 (right) expression in 786-O cells treated with vehicle control (Neg Control), scrambled siRNA (siSCR) or siRNA against LDLR (siLDLR). B-D. Rescue of ML210-induced (1 µM) cell death in 786-O neg control (B), siSCR (C) and siLDLR (D) cells with increasing concentrations of HDL, α-toc HDL, LDL and α-toc LDL. E. Rescue of Ramos, SUDHL4 and 786-O cells treated with erastin (Ramos – 4 µM; SUDHL4 – 1 µM; 786-O – 4 µM) by α-toc HDL (50 nM), α-toc LDL (50 nM) and α-toc ocHDL NPs (50 nM) in the presence or absence of BLT-1 (2 µM). Statistics: *p<0.05, **p<0.01, ***p<0.001 and ****p<0.0001 indicate statistical significance. No statistical difference (p>0.05) is noted by “ns.” See Supporting Information for more detailed statistical analyses.

To determine if α-toc is directly delivered through SR-B1, we utilized the small molecule inhibitor of SR-B1, BLT-1 (blocker of lipid transport-1)^29^. Importantly, BLT-1 blocks SR-B1-mediated molecular transport of lipoprotein cargo into the cell, including α-toc^53^, but does not inhibit lipoprotein binding to the receptor^54^. Interestingly, the addition of BLT-1 to Ramos, SUDHL4, and 786-O cells resulted in rescue of ML210-induced cell death in the absence of lipoprotein addition (Figure S4), which suggests that ML210 uptake by each of the cancer cells involves SR-B1. The SR-B1 receptor is known to be involved in the cellular delivery of drugs^55^. Therefore, to determine SR-B1’s role in the uptake of α-toc, we used erastin, which directly targets cell surface x_c_^−56^, to induce cell membrane PL oxidation and ferroptosis. As demonstrated previously, treatment of Ramos, SUDHL4 and 786-O cells with α-toc HDL, α-toc LDL and α-toc ocHDL NPs rescued erastin-induced cell death (Figure S3); however, blocking delivery of α-toc using BLT-1 resulted in cell death (**Figure 3E**). Taken together, these data indicate that SR-B1, not the LDLR, is the critical receptor for α-toc delivery from lipoproteins and that delivery of α-toc via SR-B1 is responsible for the rescue from cell membrane PL oxidation and ferroptosis.

### α-toc Delivery by HDLs via SR-B1 Rescues Differentiated Neuronal Cells from Ferroptosis

Lipid oxidative stress and ferroptosis is not limited to cancer cells. In particular, exquisite control over ferroptosis by α-toc and GPx4 has been documented in other systems, such as the hematopoietic^20^ and neurologic systems^19^. Neuronal cell death by ferroptosis can result in neurodegenerative disease. We therefore utilized an in vitro cell culture model of neurons to study the α-toc/HDL/SR-B1 axis. The neuroblastoma SH-SY5Y cell line can be differentiated in vitro by adding retinoic acid and brain derived neurotrophic factor (BDNF) with ultimate removal of serum in the cell culture media (see methods) to form non-proliferative neuron-like cells. The differentiated SH-SY5Y cells are a useful *in vitro* model to study neurodegenerative disorders. Differentiated SH-SY5Y cells experience increased reactive oxygen species, lipid peroxidation, and cell death in the presence of 1-methyl-4-phenylpyridinium (MPP), which has been used as an *in vitro* model of Parkinson’s disease^57^. Additionally, lipid peroxidation and cell death can be mitigated by addition inhibitors of ferroptosis, including ferrostatin-1 and the hydrophilic analog of Vitamin E, Trolox^57^. Accordingly, we differentiated SH-SY5Y cells into neuronal-like cells as previously described^57^. Notably, undifferentiated SH-SY5Y cells expressed SR-B1, while differentiated SH-SY5Y cells expressed little detectable SR-B1 (**Figure 4A**). This is expected, as undifferentiated SH-SY5Y cells are a highly proliferative neuroblastoma cell line while terminally differentiated SH-SY5Y display a phenotype closer to neurons. Treatment of differentiated SH-SY5Y cells with MPP increased SR-B1 expression in a time dependent manner (**Figure 4A**), and significantly increased PUFA-PL-OOH accumulation (Figure 4B). Addition of HDL with α-toc reduced PUFA-PL-OOH to baseline (**Figure 4B**) and rescued the cells from cell death (**Figure 4C**). Because the SH-SY5Y cells are grown in the absence of serum that contains lipoproteins, we attempted to add free α-toc to rescue the cells from lipid oxidation and cell death. However, this failed to reduce PUFA-PL-OOH accumulation and did not rescue the cells from MPP-induced cell death (**Figure 4B, 4C**). Presumably, this is due to the lack of lipoprotein carriers (HDL and LDL). Overall, these data demonstrate that differentiated SH-SY5Y cells in conditions of increased oxidative stress, for example with treatment by the neurotoxin MPP, increased SR-B1 expression and, thus, provide an opportunity for α-toc delivery via HDL, which rescues neuronal cell death.

**Figure 4:**
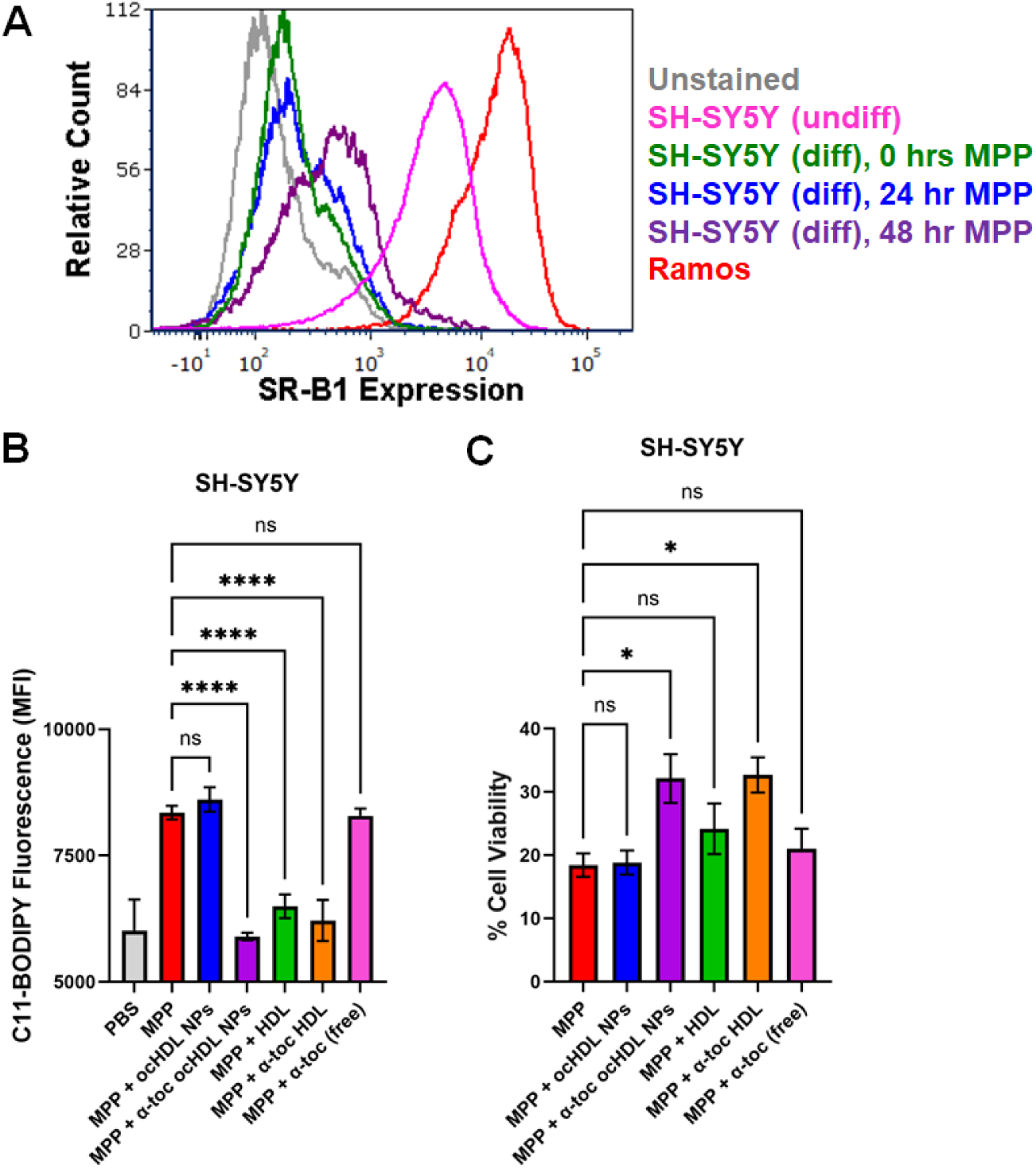
MPP-Induced Ferroptosis-Like Cell Death Rescued in Differentiated SH-SY5Y Neuronal Cells by α-toc HDLs. A. SR-B1 expression analysis by flow cytometry. Differentiated SH-SY5Y cells were treated with MPP (2.5 mM) for 0, 24 or 48 hrs prior to flow analysis. Ramos was used as an SR-B1 positive control. SH-SY5Y (undiff) = undifferentiated SH-SY5Y cells. SH-SY5Y (diff) = differentiated SH-SY5Y cells. B. SH-SY5Y cells were treated MPP (2.5 mM) in addition to 0 or 25 nM ocHDL NPs, α-toc ocHDL NPs, HDL, α-toc HDL or free α-toc (250 nM). PUFA-PL-OOH accumulation was quantified by C11 BODIPY flow cytometry 48 hrs later. Data presented as median fluorescent intensity (MFI). C. SH-SY5Y cells were treated with 2.5 mM MPP with 0 or 25 nM ocHDL NPs, α-toc ocHDL NPs, HDL, α-toc HDL or free α-toc (250 nM). Cell viability was quantified by MTS assay 48 hrs later. Statistics: *p<0.05 and ****p<0.0001 indicate statistical significance. No statistical difference (p>0.05) is noted by “ns.” See Supporting Information for more detailed statistical analyses.

## DISCUSSION

We report a physiologic α-toc/HDL/SR-B1 axis of cell membrane PL redox control in lymphoma, renal cancer and neuronal cells whereby lipoproteins, predominantly HDLs, carry α-toc, bind SR-B1, and deliver α-toc to the cell membrane to reduce lipid oxidative damage and inhibit cell death. Both natural HDL and synthetic ocHDL NP ligands bind SR-B1 and can be used to manipulate the α-toc/HDL/SR-B1 axis to control lipid oxidation and cell fate. In cancer, as suggested by our work^37,51^ and the work of others^11^, targeting multiple mechanisms through which cancer cells protect themselves from oxidative damage and ferroptosis is essential to realizing successful therapy. As in cancer cells, the redox axis is relevant to neuronal cells, as they can be rescued from lipid oxidative cell death by targeting SR-B1 with α-toc replete SR-B1 ligands, including native, functional HDLs. Prior studies show that this mechanism of neuroprotection may be highly effective in cooperation with GPx4^58,59^.

While α-toc is a well-established natural lipophilic antioxidant known to be transported in HDL and LDL, a connection has never been made between α-toc HDL or α-toc LDL, specific cell targeting, and cell membrane lipid redox balance/ ferroptosis. We have shown in these studies that by targeting SR-B1 in vitro and in vivo with lipid-poor HDL NPs we can increase cell membrane lipid oxidation and induce ferroptosis in cancer cells. Therefore, we hypothesized that the opposite may be true for native lipid (e.g., α-toc)-rich HDLs or synthetic α-toc HDL NPs that target SR-B1 and that this represents an innate, heretofore unrecognized physiologic axis that regulates cell oxidation and cell death by ferroptosis.

To validate this pathway and overcome the chemical heterogeneity challenges posed by native lipoproteins, we found that the synthetic HDL NPs with α-toc, but not the ones devoid of α-toc, also protect cells from cell membrane lipid oxidation and ferroptosis. Our findings shine a light on the perhaps underappreciated antioxidative function of native HDLs and synthetic ligands of SR-B1, which may be leveraged to develop prevention and treatment strategies for devastating diseases that result from cell membrane redox imbalance and ferroptosis.

Overall, the α-toc/HDL/SR-B1 axis of cell membrane PL redox control that we describe herein may enhance our understanding of HDL function and biology, cell membrane redox balance, and ferroptosis to provide next generation therapies for cancer, neurodegeneration, inflammation, and other diseases associated with increased lipid oxidative damage.

## MATERIALS AND METHODS

### Cell Lines and Reagents

The Ramos, SUDHL4, 786-O, U266B1 and SH-SY5Y cell lines were purchased directly from ATCC and used within 3 months of resuscitation from cryopreservation. ATCC uses short tandem repeats (STR) profiling to authenticate their cell lines. B-cell lymphoma and ccRCC cell lines were cultured in RMPI 1640 media (Corning), supplemented with 10% fetal bovine serum (FBS, Corning) and 1% PenStrep (Corning). For the lipoprotein deficient serum experiments, FBS was replaced in the culture media formulation with 10% LPDS (Kalen Biomedical). ML210 and erastin were obtained from Selleck Chemicals. BLT-1 and MPP+ (1-methyl-4-phenylpyridinium iodide) were obtained from Sigma Aldrich. The C11-BODIPY dye was obtained from ThermoFisher Scientific. HDL and LDL were obtained from Sigma Aldrich. Molar concentrations of HDL were calculated as described previously^60^. Molar concentrations of LDL were calculated by converting the mg/ mL to molarity, assuming that the entire protein concentration is comprised of apoB-100 and that there is a single apoB-100 protein per LDL.

### Loading of HDL and LDL with α-Toc

HDL and LDL were incubated with free α-toc (dissolved in DMSO) at a molar ratio of 10/ 1 α-toc/ apoA-I (HDL) or apoB-100 (LDL), for 4 hrs at room temperature on an orbital shaker (60 rpm). Following incubation, excess, unincorporated α-toc was removed by centrifugation using spin filter columns. Protein concentrations were determined using the BCA assay (Pierce BCA protein assay kit, ThermoFisher Scientific).

### Synthesis of HDL NPs

ocHDL NPs and α-toc ocHDL NPs were synthesized using a modified inhouse protocol^36^. Briefly, apolipoprotein A-I (apoA-I; MyBioSource) and sodium cholate (final concentration 19 mM; Sigma Aldrich) in water were combined with the organic core scaffold (3-(stearoyloxy)-2,2-bis[(stearoyloxy)methyl]propyl stearate; PE-S4; CAS 115-83-3; Sigma Aldrich), phospholipids (1,2-dioleoyl-sn-glycero-3-phosphocholine; DOPC; Avanti Polar Lipids) and α-toc (Sigma Aldrich) in dichloromethane (DCM). Reagents were added at a ratio of 300/ 5/ 2/ 1 DOPC/ α-toc/ apoA-I/ PE-S4, with the final resultant mixture 75% water/ 25% DCM. An emulsion was generated by successive rounds of vortexing and sonication, followed by stirring to allow the organic phase (DCM) to evaporate and the ocHDL NPs to self-assemble. The resultant nanoparticle solution was then dialyzed against 1 X PBS for 24 hours, with one buffer change at ∼12 hrs. ApoA-I concentration was calculated using a BCA assay, and the final nanoparticle concentration determined using 2.23 apoA-I/ nanoparticle.

### Characterization of Lipoproteins and HDL NPs

All lipoproteins (e.g., HDL, LDL) and ocHDL NPs were characterized by dynamic light scattering (DLS) to measure size (diameter, in nm), surface charge (zeta potential, reported in mV) and α-toc content (Vitamin E assay kit, MyBioSource, MBS2540415). DLS (3 runs per construct, with each run consisting of 12 reads of 15 sec each; sample concentration of ∼40nM in 1 X PBS) and zeta potential (3 runs per construct, with each run consisting of a minimum of 30 to a maximum of 100 measurements; sample concentration of ∼40nM in water) were measured using a Malvern Zetasizer. A UV/ Vis spectrophotometer was used to quantify the absorbance readings (533 nm) from the Vitamin E Assay Kit (N = 3 per lipoprotein/ nanoparticle).

### Quantification of Lipoprotein Receptors by Flow Cytometry

SR-B1 and LDLR expression was quantified by flow cytometry, as described previously^37^. APC-labeled SR-B1 antibody was obtained from BioLegend (#363208) and APC-labeled LDLR antibody was obtained from Abcam (AB275614). Blocking was achieved using Human TruStain FcX (BioLegend).

### Quantification of PUFA-PL-OOH Accumulation

Measurement of PUFA-PL-OOH accumulation was conducted using the C11-BODIPY fluorescent dye and flow cytometry, as described previously^37,51^. C11-BODIPY was obtained from ThermoFisher Scientific (D3861).

### Western Blot Analysis

Western blots were conducted as described previously^37,51^. Antibodies against SR-B1 (ab52629, Abcam) and GPx4 (ab125066, Abcam) were used at a 1:1000 dilution, the antibody against β-actin (#4967, Cell Signaling Technologies) was used at a 1:3000 dilution and the secondary antibodies (BioRad) were used at a 1:2000 dilution. Blots were imaged using an Azure 300 western blot imager (Azure Biosystems).

### Cell Viability Assay (MTS)

Cell viability was quantified using the MTS assay, as described previously^37,51^. Briefly, Ramos, SUDHL4 and U266B1 were plated at a density of 2 × 10^5^ cells/ mL for experiments lasting 72 hrs (cell death induced by ML210) and 1 × 10^6^ cells/ mL for experiments lasting 24 hrs (cell death induced by erastin). 786-O cells were plated at a density of 5 × 10^3^ cells/ mL for 72 hr. experiments (ML210), and 1 × 10^4^ cells/ mL for 24 hr. experiments (erastin). Promega’s CellTiter Aqueous reagent (Promega) was used to quantify percent viability, as described previously^37,51^.

### Knockdown of SR-B1 and LDLR in 786-O Cells

786-O cells were transfected with vehicle control (neg control), 50 nM scrambled siRNA (siSCR; Invitrogen), 50 nM SR-B1 siRNA (siSR-B1; Invitrogen) or 50 nM LDLR siRNA (siLDLR; Invitrogen) using Lipofectamine RNAiMAX (Invitrogen), as described previously^61^. Cells were cultured for 48 hrs prior to replating and treated with various ferroptosis inducers (ML210) and lipoproteins. Knockdown of LDLR was confirmed by flow cytometry.

### Differentiation of SH-SY5Y Cells

SH-SY5Y cells were differentiated as described previously^57^. Briefly, SH-SY5Y cells were cultured with 10 µM retinoic acid in DMEM/ Ham’s media (Corning) supplemented with 10% FBS, 2 mM glutamine and 1% PenStrep for 7 days, followed by culture in serum free DMEM/ Ham’s media with 2 mM glutamine, 1% PenStrep and 50 ng/ mL BDNF (R&D Systems) for a further 5-6 days. After differentiation, SH-SY5Y cells were plated (4 × 10^5^ cells/ mL) into collagen I coated 6 well plates (C11 BODIPY and SR-B1 flow analysis) or 96 well plates (MTS assay). Cells were allowed to adhere overnight prior to treatment with MPP+ (2.5 mM) and lipoproteins. Cells were cultured for 48 hrs prior to assaying for PUFA-PL-OOH accumulation, SR-B1 expression and viability.

### Statistical Analysis

Statistics were run in GraphPad Prism 10. N values varied between 4 to 10 for cell viability/ MTS assays (exact numbers indicated in each caption), while N = 3 for C11 BODIPY experiments. T-tests and 1-way ANOVAs, with multiple comparisons, were used for these studies. For MTS studies measuring the ability of lipoproteins to rescue small molecule induced cell death (e.g., HDL rescue of ML210 induced cell death) as well as the corresponding C11-BODIPY analysis of lipid peroxide accumulation, 1-way ANOVAs with multiple comparisons to the small molecule treatment alone were conducted. For MTS studies measuring the ability of BLT-1 to inhibit lipoprotein rescue of erastin- or ML210-induced cell death, multiple unpaired t-tests were run comparing each treatment condition without BLT1 (-BLT1) to that with BLT1 (+BLT1).

## ACKNOWLEDGEMENTS

This work was supported by a V Foundation grant (T2022-018, CST). We thank the Robert H. Lurie Comprehensive Cancer Center of Northwestern University in Chicago, IL for the use of the Flow Cytometry Core Facility, which provided access to the BD Fortessa II cytometer. The Lurie Cancer Center is supported in part by an NCI Cancer Center Support Grant #P30CA060553. The Azure 300 imaging system was used in the Analytical bioNanoTechnology Equipment Core Facility of the Simpson Querrey Institute for BioNanotechnology at Northwestern University. ANTEC receives partial support from the Soft Hybrid Nanotechnology Experimental (ShyNE) Resources (NSF ECCS-2025633) and Feinberg School of Medicine, Northwestern University.

## SUPPORTING INFORMATION

### Figure Statistics

**Figure 1:** A-C. N = 4 per group. 1-way ANOVA tests (F, degrees of freedom, p value, R^2^), with multiple comparisons between 0 nM and all other groups. Ramos HDL: (58.11, 3, <0.0001, 0.9561); 0nM vs. 10nM = 0.9799; 0nM vs. 25nM = 0.1981; ****0nM vs. 50nM < 0.0001. Ramos α-toc HDL: (259.3, 3, <0.0001, 0.9861); 0.3465; ****<0.0001; ****<0.0001. Ramos LDL: (12.54, 3, 0.0022, 0.8246); 0.1563; 0.1563; ***0.0008. Ramos α-toc LDL: (63.21, 3, <0.0001, 0.9595); 0.6427; 0.8861; ****<0.0001. Ramos ocHDL NP: (2.779, 3, 0.0911, 0.4312); 0.9893; 0.0987; 0.8898. Ramos α-toc ocHDL NP: (403.9, 3, <0.0001, 0.9910); ****<0.0001; ****<0.0001; ****<0.0001. SUDHL4 HDL: (14.70, 3, 0.0013, 0.8464); 0nM vs. 10nM = 0.9809; 0nM vs. 25nM = 0.9238; **0nM vs. 50nM = 0.0012. SUDHL4 α-toc HDL: (1405, 3, <0.0001, 0.9981); *0.0126; ****<0.0001; ****<0.0001. SUDHL4 LDL: (5.597, 3, 0.0230, 0.6773); 0.4972; 0.2446; **0.0100. SUDHL4 α-toc LDL: (124.1, 3, <0.0001, 0.9713); 0.9999; 0.6477; ****<0.0001. SUDHL4 ocHDL NP: (0.1937, 3, 0.8985, 0.05019); 0.9982; 0.9561; 0.8371. SUDHL4 α-toc ocHDL NP: (467.2, 3, <0.0001, 0.9922); ****<0.0001; ****<0.0001; ****<0.0001. 786-O HDL: (94.85, 3, <0.0001, 0.9727); 0nM vs. 10nM = 0.8088; 0nM vs. 25nM = 0.2012; ****0nM vs. 50nM <0.0001. 786-O α-toc HDL: (308.4, 3, <0.0001, 0.9914); 0.0908; ****<0.0001; ****<0.0001. 786-O LDL: (2.946, 3, 0.0801, 0.4455); 0.0659; 0.1260; 0.6928. 786-O α-toc LDL: (158.8, 3, <0.0001, 0.9774); 0.6583; ****<0.0001; ****<0.0001. 786-O ocHDL NP: (10.66, 3, 0.0014, 0.7441); **0.0027; **0.0045; **0.0058. 786-O α-toc ocHDL NP: (784.7, 3, <0.0001, 0.9953); ****<0.0001; ****<0.0001; ****<0.0001. D. N = 3 per group. 1-way ANOVA tests (F, degrees of freedom, p value, R^2^), with multiple comparisons between 0 nM and all other groups. HDL: (7.262, 2, 0.0250, 0.7076); *0nM vs. 10nM = 0.0301; *0nM vs. 50nM = 0.0278. α-toc HDL: (21.15, 2, 0.0019, 0.8758); **0.0022; **0.0026. LDL: (0.9997, 2, 0.4220, 0.2500); 0.9179; 0.3550. α-toc LDL: (10.54, 2, 0.0109, 0.7784); *0.0308; **0.0079. ocHDL NP: (0.9049, 2, 0.4534, 0.2317); 0nM vs. 10nM = 0.4526; 0nM vs. 50nM = 0.9998. α-toc ocHDL NP: (18.61, 2, 0.0027, 0.8612); **0.0049; **0.0025.

**Figure 2:** N = 4 per group. 1-way ANOVA tests (F, degrees of freedom, p value, R^2^), with multiple comparisons between 0 nM and all other groups. Ramos HDL: (4.771, 3, 0.0115, 0.4171); 0nM vs. 10nM = 0.4809; 0nM vs. 25nM = 0.0758; **0nM vs. 50nM = 0.0075. Ramos α-toc HDL: (246.2, 3, <0.0001, 0.9736); ****<0.0001; ****<0.0001; ****<0.0001. Ramos LDL: (20.36, 3, <0.0001, 0.7533); ***0.0009; ****<0.0001; ****<0.0001. Ramos α-toc LDL: (32.21, 3, <0.0001, 0.8285); **0.0067; ****<0.0001; ****<0.0001. Ramos ocHDL NP: (6.515, 3, 0.003, 0.4943); 0.0578; **0.0032; *0.0359. Ramos α-toc ocHDL NP: (193.9, 3, <0.0001, 0.9668); ****<0.0001; ****<0.0001; ****<0.0001. SUDHL4 HDL: (10.76, 3, 0.0002, 0.6174); 0nM vs. 10nM = 0.1866; **0nM vs. 25nM = 0.0053; ***0nM vs. 50nM = 0.0001. SUDHL4 α-toc HDL: (200.7, 3, <0.0001, 0.9678); ****<0.0001; ****<0.0001; ****<0.0001. SUDHL4 LDL: (5.008, 3, 0.0095, 0.4289); *0.0332; 0.5945; 0.3501. SUDHL4 α-toc LDL: (30.08, 3, <0.0001, 0.8186); **0.0020; **0.0015; ****<0.0001. SUDHL4 ocHDL NP: (1.236, 3, 0.3228, 0.1564); 0.9855; 0.2607; 0.9977. SUDHL4 α-toc ocHDL NP: (223.9, 3, <0.0001, 0.9711); ****<0.0001; ****<0.0001; ****<0.0001. 786-O HDL: (83.52, 3, <0.0001, 0.9261); 0nM vs. 10nM = 0.7446; 0nM vs. 25nM = 0.8953; ****0nM vs. 50nM <0.0001. 786-O α-toc HDL: (2169, 3, <0.0001, 0.9969); ****<0.0001; ****<0.0001; ****<0.0001. 786-O LDL: (3.509, 3, 0.0342, 0.3448); 0.2381; 0.9134; 0.1278. 786-O α-toc LDL: (3.507, 3, 0.0343, 0.3447); 0.0579; 0.9956; 0.4734. 786-O ocHDL NP: (8.477, 3, 0.008, 0.5598); **0.0028; **0.0073; **0.0097. 786-O α-toc ocHDL NP: (587.1, 3, <0.0001, 0.9888); ****<0.0001; ****<0.0001; ****<0.0001.

**Figure 3:** B-D. N = 4 per group. 1-way ANOVA tests (F, degrees of freedom, p value, R^2^), with multiple comparisons between 0 nM and all other groups. Neg Control HDL: (29.20, 3, <0.0001, 0.8884); 0nM vs. 10nM = 0.9804; 0nM vs. 25nM = 0.1164; ****0nM vs. 50nM < 0.0001. Neg Control α-toc HDL: (1309, 3, <0.0001, 0.9972); ****<0.0001; ****<0.0001; ****<0.0001. Neg Control LDL: (2.400, 3, 0.1233, 0.3956); 0.9497; 0.6321; 0.1154. Neg Control α-toc LDL: (58.79, 3, <0.0001, 0.9413); >0.9999; 0.2094; ****<0.0001. siSCR HDL: (12.19, 3, 0.0008, 0.7688); 0nM vs. 10nM > 0.9999; 0nM vs. 25nM = 0.9940; ***0nM vs. 50nM = 0.0005. siSCR α-toc HDL: (414.2, 3, <0.0001, 0.9911); ****<0.0001; ****<0.0001; ****<0.0001. siSCR LDL: (11.90, 3, 0.0009, 0.7644); 0.6478; 0.0907; ***0.0003. siSCR α-toc LDL: (291.7, 3, <0.0001, 0.9876); 0.7402; ***0.0002; ****<0.0001. siLDLR HDL: (110.2, 3, <0.0001, 0.9678); 0nM vs. 10nM = 0.2448; **0nM vs. 25nM = 0.0049; ****0nM vs. 50nM <0.0001. siLDLR α-toc HDL: (400.6, 3, <0.0001, 0.9909); ****<0.0001; ****<0.0001; ****<0.0001. siLDLR LDL: (37.29, 3, <0.0001, 0.9105); 0.1119; ***0.0002; ****<0.0001. siLDLR α-toc LDL: (202.2, 3, <0.0001, 0.9822); ****<0.0001; ****<0.0001; ****<0.0001. E. N = 5 per group. Multiple unpaired t tests were run, comparing -BLT1 and +BLT1 groups for each treatment combination (e.g., PBS, erastin). Ramos: PBS = 0.187515; Erastin = 0.255202; Erastin + α-toc HDL <0.000001; Erastin + α-toc LDL = 0.000008; Erastin + α-toc ocHDL NP = 0.000005. SUDHL4: PBS = 0.456104; Erastin = 0.574740; Erastin + α-toc HDL = 0.000079; Erastin + α-toc LDL = 0.000096; Erastin + α-toc ocHDL NP = 0.000042. 786-O: PBS = 0.365859; Erastin = 0.176284; Erastin + α-toc HDL = 0.000078; Erastin + α-toc LDL = 0.049870; Erastin + α-toc ocHDL NP = 0.000002.

**Figure 4:** B. N = 3 per group. 1-way ANOVA tests (F = 40.00, degrees of freedom = 6, p value < 0.0001, R^2^ = 0.9486), with multiple comparisons between MPP and all other groups. MPP vs. MPP + ocHDL NPs = 0.8395; ****MPP vs. MPP + α-toc ocHDL NPs < 0.0001; ****MPP vs. MPP + HDL < 0.0001; ****MPP vs. MPP + α-toc HDL < 0.0001; MPP vs. MPP + α-toc (free) > 0.9999; MPP vs. PBS < 0.0001. C. N = 6 per group. 1-way ANOVA tests (F = 4.475, degrees of freedom = 5, p value = 0.0036, R^2^ = 0.4272), with multiple comparisons between MPP and all other groups. MPP vs. MPP + ocHDL NPs > 0.9999; *MPP vs. MPP + α-toc ocHDL NPs = 0.0140; *MPP vs. MPP + HDL = 0.5386; *MPP vs. MPP + α-toc HDL = 0.0101; MPP vs. MPP + α-toc (free) = 0.9579.

**Table S1:**
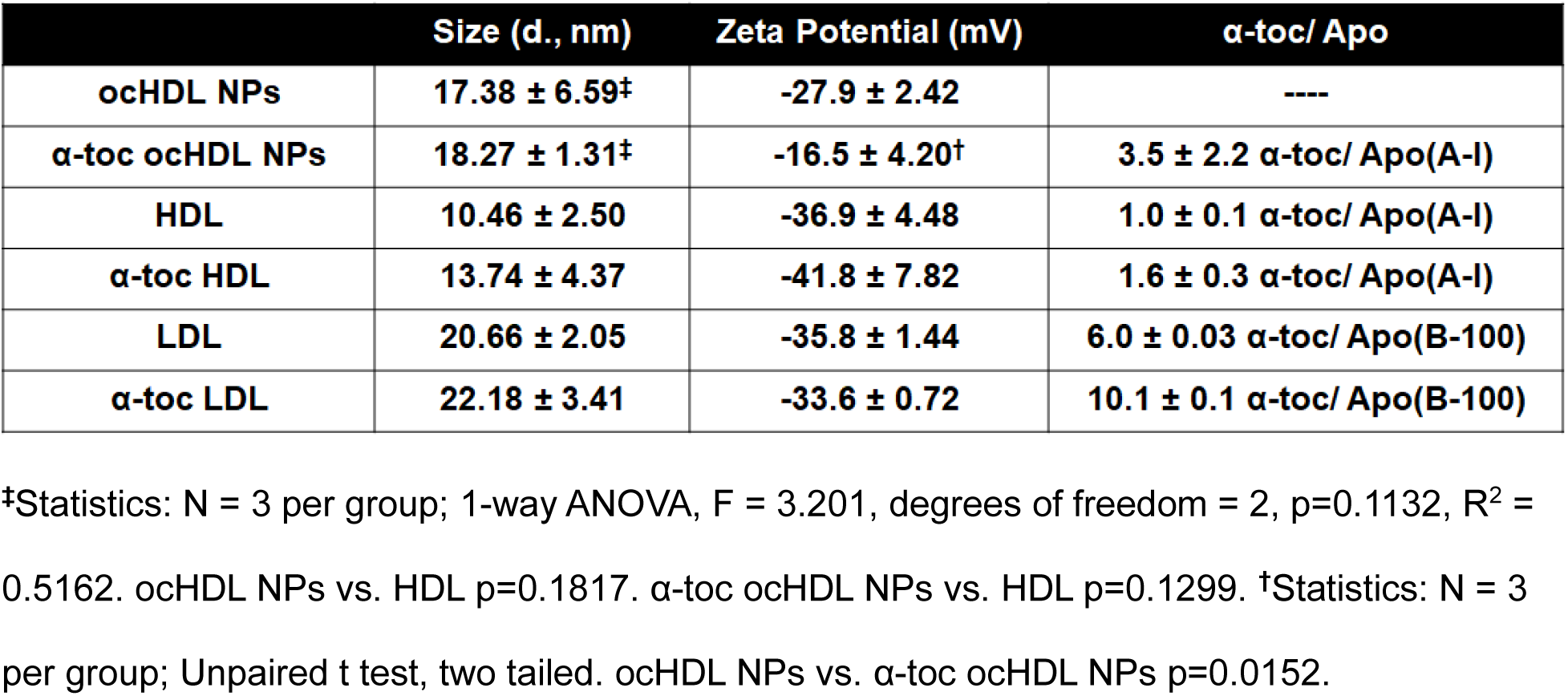
Characterization of HDL and LDL Nanoparticles.

| | Size (d., nm) | Zeta Potential (mV) | α-toc/ Apo |
| --- | --- | --- | --- |
| ocHDL NPs | $17.38 \pm 6.59^\ddagger$ | $-27.9 \pm 2.42$ | ---- |
| α-toc ocHDL NPs | $18.27 \pm 1.31^\ddagger$ | $-16.5 \pm 4.20^\dagger$ | $3.5 \pm 2.2$ α-toc/ Apo(A-I) |
| HDL | $10.46 \pm 2.50$ | $-36.9 \pm 4.48$ | $1.0 \pm 0.1$ α-toc/ Apo(A-I) |
| α-toc HDL | $13.74 \pm 4.37$ | $-41.8 \pm 7.82$ | $1.6 \pm 0.3$ α-toc/ Apo(A-I) |
| LDL | $20.66 \pm 2.05$ | $-35.8 \pm 1.44$ | $6.0 \pm 0.03$ α-toc/ Apo(B-100) |
| α-toc LDL | $22.18 \pm 3.41$ | $-33.6 \pm 0.72$ | $10.1 \pm 0.1$ α-toc/ Apo(B-100) |

$^\ddagger$ Statistics: N = 3 per group; 1-way ANOVA, $F = 3.201$ , degrees of freedom = 2, $p=0.1132$ , $R^2 = 0.5162$ . ocHDL NPs vs. HDL $p=0.1817$ . α-toc ocHDL NPs vs. HDL $p=0.1299$ . $^\dagger$ Statistics: N = 3 per group; Unpaired t test, two tailed. ocHDL NPs vs. α-toc ocHDL NPs $p=0.0152$ .

**Figure S1:**
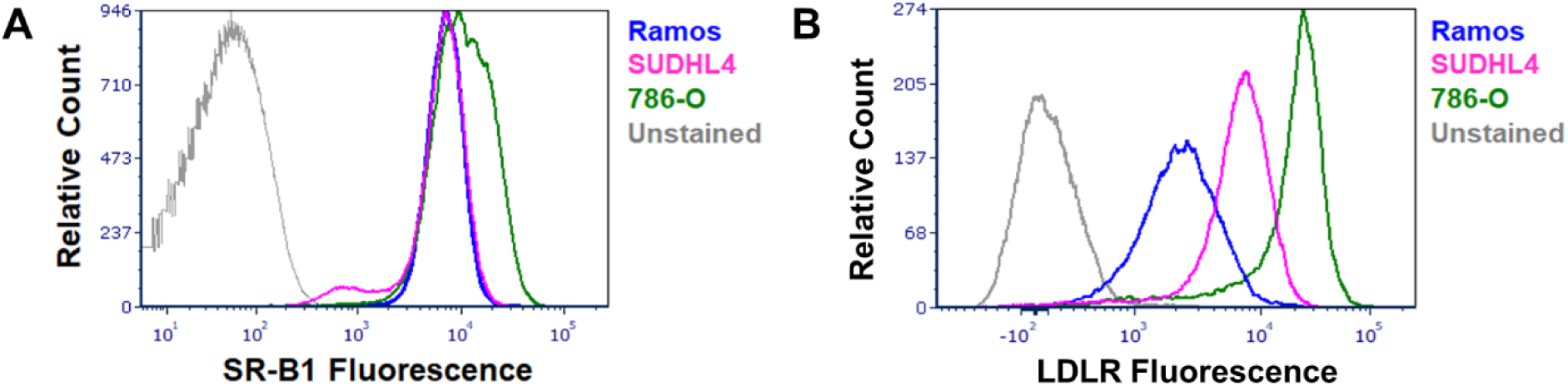
SR-B1 and LDLR Expression. Flow cytometry analysis of SR-B1 (A) and LDLR (B) expression in the lymphoma cell lines Ramos and SUDHL4 and the renal cell carcinoma cell line 786-O.

**Figure S2:**
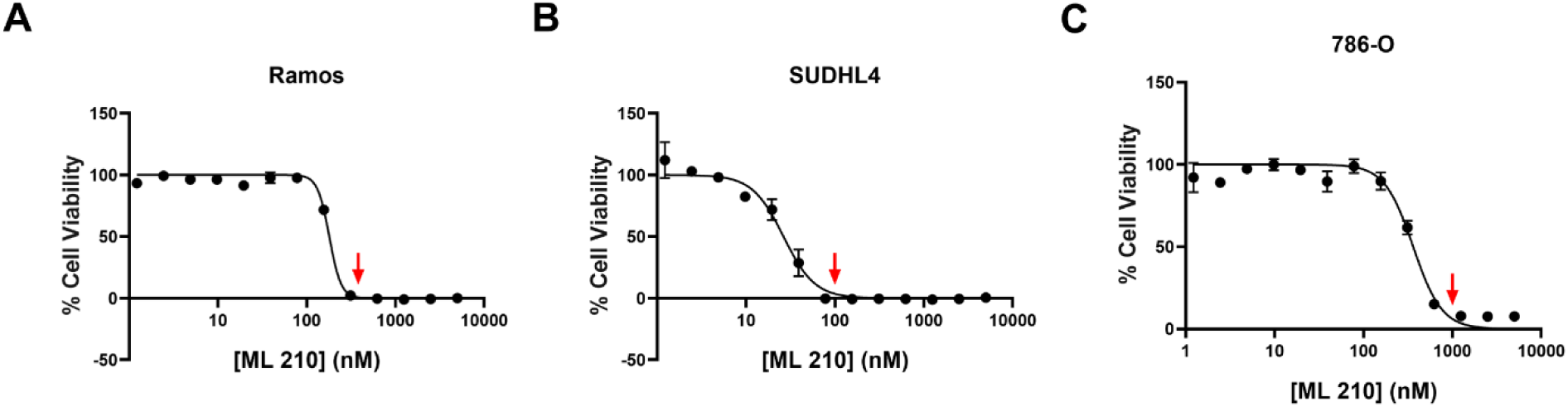
ML210 Induces Cell Death in DLBCL and ccRCC Cell Lines. A-C. ML210 kill curves for Ramos (A), SUDHL4 (B) and 786-O (C) cells. Red arrow indicates concentration used in the α-toc lipoprotein rescue experiments detailed in Figures 1 and 2.

**Figure S3:**
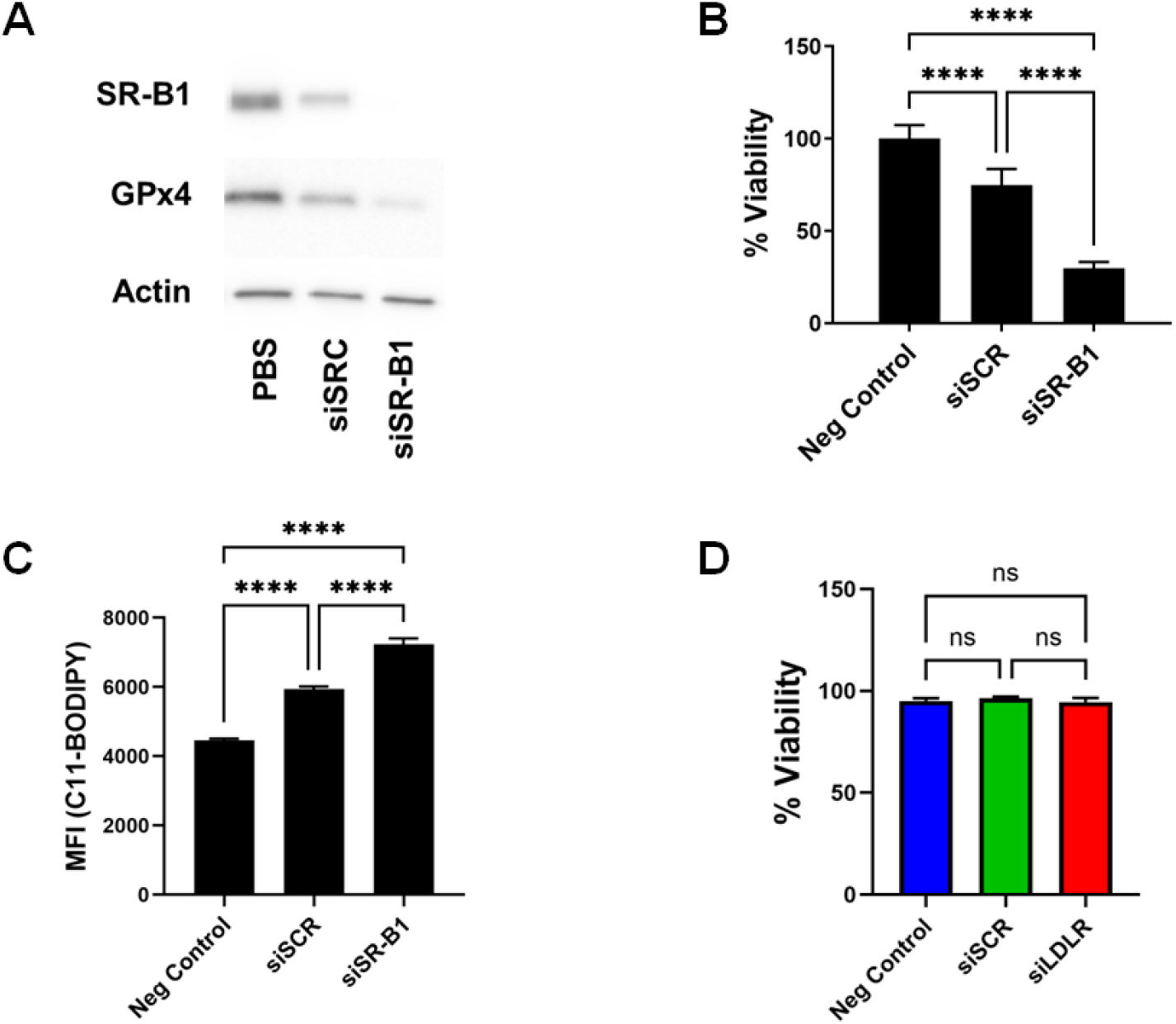
Knockdown of SR-B1 Induces Ferroptosis in 786-O Cells. A. siRNA mediated knockdown of SR-B1 (siSR-B1) reduced both SR-B1 and GPx4 expression in 786-O cells, while a lesser change in expression of either protein were observed in the scrambled control siRNA (siSCR). B. siSR-B1 treatment reduced 786-O cell viability at 5 days post knockdown. Statistics: N = 10 per group; 1-way ANOVA, F = 267.3, degrees of freedom = 2, p<0.0001, R^2^ = 0.9519. ****p<0.0001. C. PUFA-PL-OOH accumulation increases with knockdown of SR-B1. Statistics: N = 5 per group; 1-way ANOVA, F = 155.2, degrees of freedom = 2, p<0.0001, R^2^ = 0.9628. ****p<0.0001. D siRNA mediated knockdown of LDLR did not alter the viability of 786-O cells. Statistics: N = 3 per group; 1-way ANOVA, F = 0.9286, degrees of freedom = 3, p=0.4854, R2 = 0.3824. P values: Neg control vs. siSCR = 0.6350; Neg control vs. siLDLR = 0.9437; siSCR vs. siLDLR = 0.4815.

**Figure S4:**
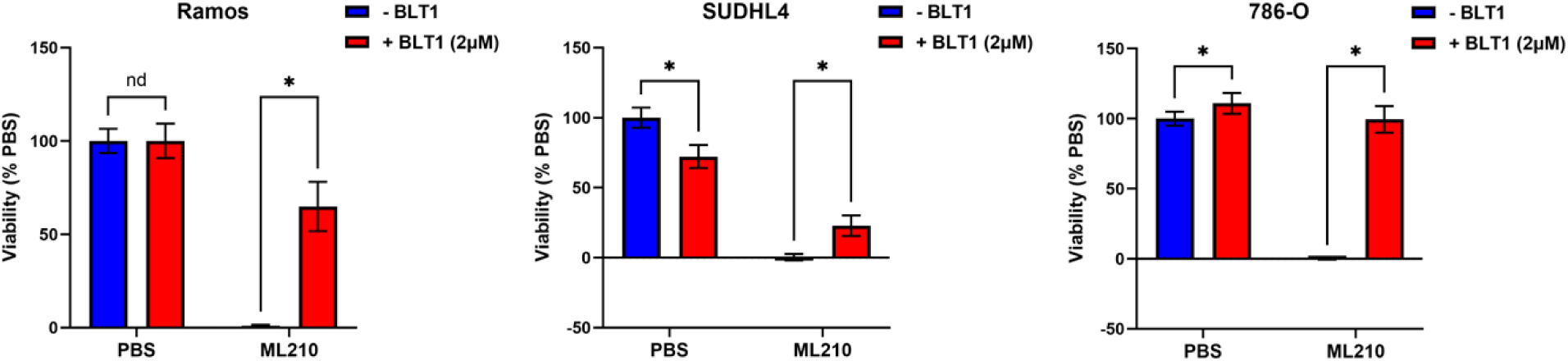
BLT-1 Treatment Rescues Cells from ML210-Induced Cell Death. Ramos, SUDHL4 and 786-O cells were treated with PBS or ML210 (250 nM for Ramos; 100 nM for SUDHL4; 1 µM for 786-O) in the presence or absence of BLT-1 (1 µM). Viability was quantified by MTS assay. Data are presented relative to PBS control without BLT-1 (-BLT-1). Statistics: N = 5 per group. Multiple unpaired t-tests (-BLT1 vs. + BLT1; 2 tests performed per cell line). Ramos p values: PBS = 0.987064, ML210 < 0.000001. SUDHL4 p values: PBS = 0.000097, ML210 = 0.000032. 786-O p values: PBS = 0.013589, ML210 < 0.000001.

## Notes

### Competing Interest Statement

The authors have declared no competing interest.

